# An integrated scalable process for adherent cultivated meat production: From proliferative cell selection to safety-verified product development

**DOI:** 10.64898/2026.01.08.698307

**Authors:** Hiroaki Hatano, Ibuki Kokido, Keita Tanaka, Satoshi Inoue, Masanobu Kowaka, Takahiro Sunaga, Naomi Nakamura, Misaki Sawada, Hiroaki Kondo, Kazuhiro Kunimasa, Ikko Kawashima

**Affiliations:** IntegriCulture Inc., Shonan Health Innovation Park (Shonan iPark), 2-26-1 Muraokahigashi, Fujisawa, Kanagawa, Japan; Hamano Products Co., Ltd., 4-39-7 Yahiro, Sumida-ku, Tokyo 131-0041, Japan

**Keywords:** Cellular agriculture, Cultivated meat, Cell-based food;Packed-bed bioreactor, Bioprocess development, Adherent cell cultivation

## Abstract

Cultivated meat can contribute to the global food supply; therefore, establishing efficient production processes is an urgent task to meet the growing demand for sustainable protein. Although cell culture technology has historically focused on suspension cells in the pharmaceutical and, subsequently, cultivated meat industries, the development of efficient processes for most adhesion-dependent cells has lagged, with limited examples reported to date. Therefore, this study used primary duck liver-derived adherent cells to develop and evaluate an integrated three-step meat production process involving pre-culture to select highly proliferative cells, packed-bed bioreactor expansion for mass production, and final processing, including packaging and heating.

The established process allowed the efficient growth of selected cell populations while maintaining their proliferative characteristics. Moreover, the developed product met the microbiological and heavy metal safety criteria. Comprehensive compositional analysis (nutritional, amino acid, and fatty acid profiles) revealed that the product exhibited a protein profile distinct from that of conventional duck liver paste, along with a unique lower-fat signature. As productivity is dependent on cell doubling time, culture duration and monthly production were estimated for cells from various animal species.

Overall, this study established a practical integrated production process for cultivated meat using adherent cells, providing a technological foundation for cellular agriculture applicable to diverse cell types and useful for future food supply diversification.

**GraphicalAbstract:** 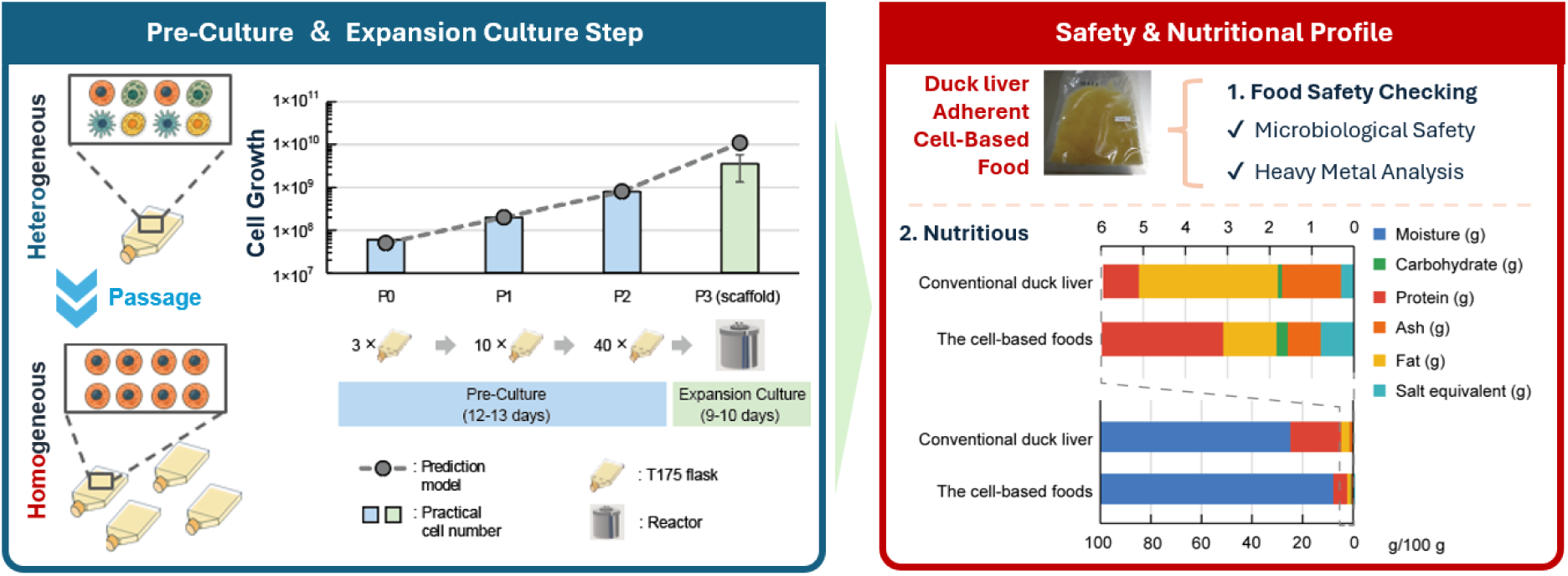

## 1. Introduction

With the growing global population, associated increase in protein demand, and increasing awareness of sustainable food production, cultivated meat production is rapidly gaining importance as an innovative approach for future food supply (Barzee et al., 2022; Bhat et al., 2014; Goodwin and Shoulders, 2013; Post, 2012; Stephens et al., 2018; Tuomisto and Teixeira de Mattos, 2011). This technology, an environment-independent production process, confers inherent resilience to climate change (Jahir et al., 2023; Reiss et al., 2021). It also improves resource utilization efficiency and reduces the environmental burden (FAO, 2024). Moreover, it has potential benefits for food security (Chriki and Hocquette, 2020). Therefore, cultivated meat is a sustainable next-generation protein source.

Currently, the cultivated meat industry is in its nascent stages, with food approval process becoming clear and products being approved by regulatory authorities and introduced to markets in some countries and regions (Mackenzi et al., 2024). These trends suggest that cultivated meat is transitioning from laboratory research to industrial applications, raising expectations across the industry (Hocquette, 2016; Nobre, 2022; Specht, 2020). However, this field is still in the early stages of development, with only a few products having received manufacturing and sales approval worldwide to date (Kirsch et al., 2023). This limitation, compounded by the historical optimization of cell culture technology primarily for pharmaceutical applications, indicates that robust and validated production processes for efficiently culturing and commercializing adhesion-dependent cells on a large scale remain scarce (Humbird, 2021; Specht et al., 2018).

The fundamental challenge is rooted in the historical evolution of cell culture technology. Large-scale processes were initially optimized for pharmaceutical manufacturing, specifically for suspension-adapted cells (e.g., Chinese hamster ovary cells for antibody drug production and Vero cells for vaccine production) (Clincke et al., 2013; Kiesslich and Kamen, 2020). Consequently, the industry possesses extensive knowledge and equipment designed for floating and suspension cell cultures (Nienow, 2006; Shukla and Gottschalk, 2013; Yao and Asayama, 2017). Consistently, the cultivated meat sector has also prioritized suspension-adapted cells to leverage existing tools (Humbird, 2021; Specht et al., 2018).

Notably, “food” requires a different approach compared to pharmaceuticals. Unlike uniform pharmaceuticals, the appeal of food lies in diversity, particularly in the unique flavors and textures of various animal species (e.g., beef, pork, and seafood) and tissues (e.g., muscle, fat, and liver) (Jayasinghe et al., 2025; Köster, 2009; Fu et al., 2022; Purslow, 2005). Use of specific cells derived from different animal species and tissues is necessary to reproduce the characteristics of these diverse meats via cell culture. Most of these cells are adhesion-dependent, requiring an attachment substrate or scaffold similar to their in vivo environment (Flores-Jiménez et al., 2023; Langer and Vacanti, 1993). Therefore, to establish cultivated meat as a diverse cultural option, the industry must move beyond suspension systems and establish efficient scalable production techniques for adhesion-dependent cells. Although existing adherent cell culture methods, such as multilayer flasks, roller bottles, and microcarrier systems, are useful at the laboratory level, breakthroughs are necessary to achieve ease of scale-up, simplify operations, reduce costs, and ensure the availability of food-grade materials suitable for food production (Hubalek et al., 2022; O’Neill et al., 2021).

In this study, we aimed to establish a scalable production process for adherent cells using duck liver as a model. We developed an end-to-end workflow to select highly proliferative cells and efficiently expand them in a packed-bed bioreactor. This approach facilitated the production of biomass meeting strict food safety standards and nutritional requirements, demonstrating the feasibility of the established system for real-world food applications.

## 2. Materials and Methods

### 2.1 Cell culture and pre-culture

Fertilized duck eggs used as the source of primary liver-derived cells were obtained from a certified egg production farm (Shiina Artificial Hatchery Co., Ltd., Chiba, Japan). Control (conventional duck liver) samples were also sourced from the same supplier to ensure a consistent genetic background, as detailed in section 2.5.

Primary duck liver-derived cells were used as the experimental model. In the pre-culture step, proprietary food-grade reagents from IntegriCulture Inc. (Kanagawa, Japan) were used. T175 flasks (VTC-F175P; VIOLAMO; AS ONE Corporation; Cat: 2-8589-15) were coated with iCoater (Hatano et al., 2025a). The cells were cultured in I-MEM 1.0 (Dulbecco’s modified Eagle’s medium equivalent; previously developed by our group (Kanayama et al., 2022)) or Dulbecco’s modified Eagle’s medium supplemented with IntegriCulture Serum-Replacement (fetal bovine serum [FBS] alternative) and maintained in a humidified incubator at 37°C with 5% CO_2_. Cell counts and viability throughout the culture period were determined using a NucleoCounter NC-202 (ChemoMetec A/S; distributed by MS Techno System, Japan) with Via2-Cassette (Cat: 941-0024; ChemoMetec A/S; distributed by MS Techno System) containing acridine orange and 4′,6-diamidino-2-phenylindole fluorescent dyes, according to the manufacturer’s instructions. For each of the two passages in the pre-culture regimen for cell expansion, the cells were initially seeded at 25% confluence (5 × 10^6^ cells/T175 flask) based on the cell count and cultured until an approximate 4-fold expansion to reach approximately 100% confluence (20 × 10^6^ cells/T175 flask), at which point they were detached using iDisper (Integriculture Inc.) (Hatano et al., 2025b). The number of passages was selected based on our previous findings that a stable and highly proliferative cell population can be effectively selected by the second passage (Hatano et al., 2025b). To determine the cell doubling time, the cells were typically cultured and analyzed while growing up to approximately 70% confluence.

### 2.2 Expansion, harvesting, packaging, and heating

For the expansion culture step, an in-house custom-designed packed-bed bioreactor with a gelatin-based scaffold (Hatano et al., 2024) was used. This custom design was inspired by the need to address challenges unresolved by existing systems. Specifically, the central rotating body of the bioreactor facilitated gentle agitation sufficient for uniform nutrient and oxygen supply to adherent cells fixed on a stationary scaffold, while the mesh cargo securely housed the scaffold. This system was specifically engineered for high-density cultivation of adherent cells, facilitating efficient cell proliferation in a three-dimensional environment. Following cultivation, the cell paste, along with the associated scaffold material, was harvested.

The harvested cultured cell paste was vacuum-packed into food-grade pouches, with each pouch containing approximately 50 g of the product. To pasteurize the product, these sealed pouches were subjected to heat treatment in a precisely controlled water bath at 75°C for 10 min, followed immediately by rapid cooling in an ice water bath. After cooling, all packages were inspected for metallic foreign objects using an industrial metal detector. The final packaged product was stored frozen at –20°C until further characterization, sensory evaluation, or other applications.

### 2.3 Fluorescence microscopy

To visualize cell viability and growth on the scaffold, the samples were stained using fluorescent dyes. The scaffold was washed with I-MEM and incubated with the -Cellstain-Calcein-AM solution (Cat: 341-07901; Dojindo) for live cells or -Cellstain-PI solution (Cat: 341-07881; Dojindo) for dead cells, along with Hoechst 33342 for nuclear staining. Images were captured using a fluorescence microscope (BZ-X810; Keyence).

### 2.4 Food safety analysis

The final processed cell paste was characterized for its suitability as a food product. All safety and nutritional analyses were conducted by an accredited third-party analytical laboratory (Food Analysis Technology Center SUNATEC, Japan). Microbiological safety assessments were conducted according to standardized food safety testing protocols (e.g., standard agar medium method for total viable counts). The tests included those shown in **Fig. 5**: *Salmonella* spp., coliforms, anaerobic spore-forming bacteria, molds, and yeasts. Heavy metal analyses of arsenic (As), lead (Pb), cadmium (Cd), and total mercury (Hg) were conducted via inductively coupled plasma mass spectrometry.

### 2.5 Nutritional analysis

Comprehensive nutritional analysis of the final cell-based product (*n* = 3–6 lots) was performed by SUNATEC. All analyses were conducted according to the official Japanese Food Labeling Standards (Notification No. 139; March 30, 2015). Specifically, proximate composition (**Fig. 6b**) was analyzed via vacuum oven drying (moisture), combustion (proteins; N factor: 6.25), acid hydrolysis (lipids), and direct ashing (ash). Energy (kcal; **Fig. 6a**) was calculated from these components. Amino acid (**Fig. 6c**) and fatty acid (**Fig. 6d**) profiles were also determined by SUNATEC using standard analytical methods.

The conventional duck liver shown in **Fig. 6** was prepared in-house to ensure precise comparison. Control (liver) samples were obtained from adult ducks from the same supplier as that for the fertilized eggs. The obtained liver samples were isolated, washed twice with MilliQ water, and processed into a paste using a mixer. This control liver paste was subjected to the same downstream processing, including vacuum packaging, heat sterilization (75°C for 10 min), rapid cooling, and frozen storage (−20°C), as the cultivated meat (detailed in **Fig. 5**). The resulting control paste was analyzed by SUNATEC under identical conditions.

### 2.6 Statistical analyses

Pearson’s correlation coefficient (*r*) was calculated to evaluate the linear associations between variables such as comprehensive gene expression datasets.

## 3. Results

### 3.1 Establishment of cultivated meat production processes for adherent cells using a packed-bed bioreactor

In this study, we developed a scalable production process using cultured duck liver-derived cells. The process was quantitatively managed based on the defined cell number metrics at each critical stage. It consisted of (1) a pre-culture step to selectively amplify a cell population with high proliferation potential and ensure sufficient cell numbers to seed the bioreactor, (2) an expansion step to maximize production using a three-dimensional high-density cell culture, and (3) a processing and packaging step to ensure the food safety of the cultured cell paste (**Fig. 1a**).

**Fig. 1.**
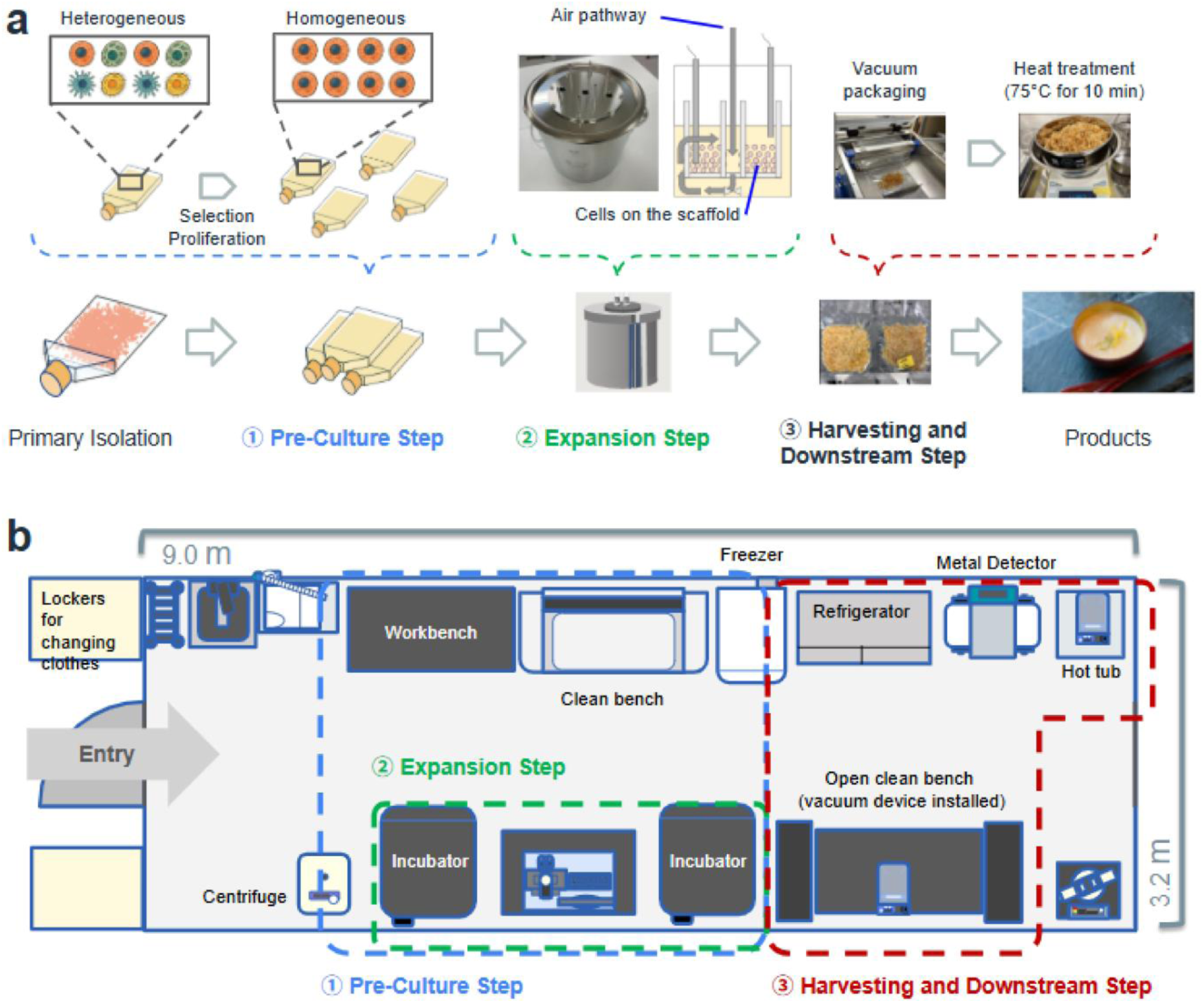
A standardized end-to-end production process for adherent cell-based foods. (a) Schematic of the integrated production workflow. The process comprised three main steps: (1) A pre-culture step for selective amplification of high-proliferation duck liver-derived cells, (2) an expansion step involving three-dimensional high-density culture in a packed-bed bioreactor, and (3) a processing and packaging step, including vacuum packaging and heat sterilization (75 °C for 10 min), for the cultured cell paste to ensure food safety. (b) Layout of the pilot-scale production unit. The diagram illustrates the 28.8-m² clean room facility with two bioreactors and two incubators, showing the operational arrangement.

In this production flow, the initial cell input was set at a standard of 0.9–1.9 × 10⁷ cells (data not shown). In the pre-culture step, the cells were seeded at 25% confluence in T175 flasks, cultured until they reached 100% confluence, and passaged twice. The passaging process selectively amplified the cell populations with stable and high proliferation rates, thereby generating the required number of cells for the subsequent expansion step. To seed the bioreactor, a sufficient quantity of 100% confluent T175 flasks, defined as 40 flasks (approximately 8 × 10^8^ cells), was prepared by P2 to provide the required number of cells for the subsequent expansion step.

For the expansion step, our in-house packed-bed bioreactor equipped with a food-grade cell-adherent scaffold was used. The scaffold functioned as a filler, providing a three-dimensional culture environment for adherent cells and enhancing cell growth beyond conventional two-dimensional culture characteristics (Hatano et al., 2024). This configuration enabled mass culture with high spatial efficiency, while preserving the cell growth characteristics observed in planar cultures. The manufacturing acceptance threshold for final cell recovery from the bioreactor was set at 3.6 × 10^9^ cells/batch, a target derived from the minimum yield required to produce a viable 50-g food prototype.

After cultivation, the cell paste, along with the scaffold, was harvested from the bioreactor and subjected to packaging and heating, following conventional food manufacturing procedures. Specifically, the cultured cell food paste was vacuum-packed at 50 g per bag, sterilized via heating in a 75°C hot water bath for 10 min, followed by rapid cooling in ice water. Subsequently, the paste was inspected for foreign objects using a metal detector and stored frozen at –20°C. The integrated production line established in this study was set up in a 28.8-m^2^ clean room, which currently houses two bioreactors on a trial basis, for food production (**Fig. 1b**).

### 3.2 Detailed workflow of the integrated production process

The scalable production process developed in this study is illustrated in **Fig. 2**. The integrated workflow consisted of two main steps: A preculture step and an expansion culture step. The pre-culture step (**Fig. 2a**) was designed to selectively amplify a cell population with high proliferation potential. This was achieved by obtaining primary cells through two passages in T175 flasks. This standardized procedure ensured the generation of a homogeneous cell population meeting the quantitative metrics required for seeding the bioreactor. Subsequently, the process transitioned to the expansion culture step (**Fig. 2b**). This step began with a “preparation phase,” where the custom packed-bed bioreactor (its structure is shown in **Fig. 3**) was prepared with scaffolds and medium before the introduction of the cell population from the pre-culture step. This was followed by the “cultivation phase” for 9–10 days. After this phase, the cell biomass was harvested and subjected to downstream processing, including vacuum packaging, heat pasteurization, and inspection for metallic foreign objects, before being stored as the final product.

**Fig. 2.**
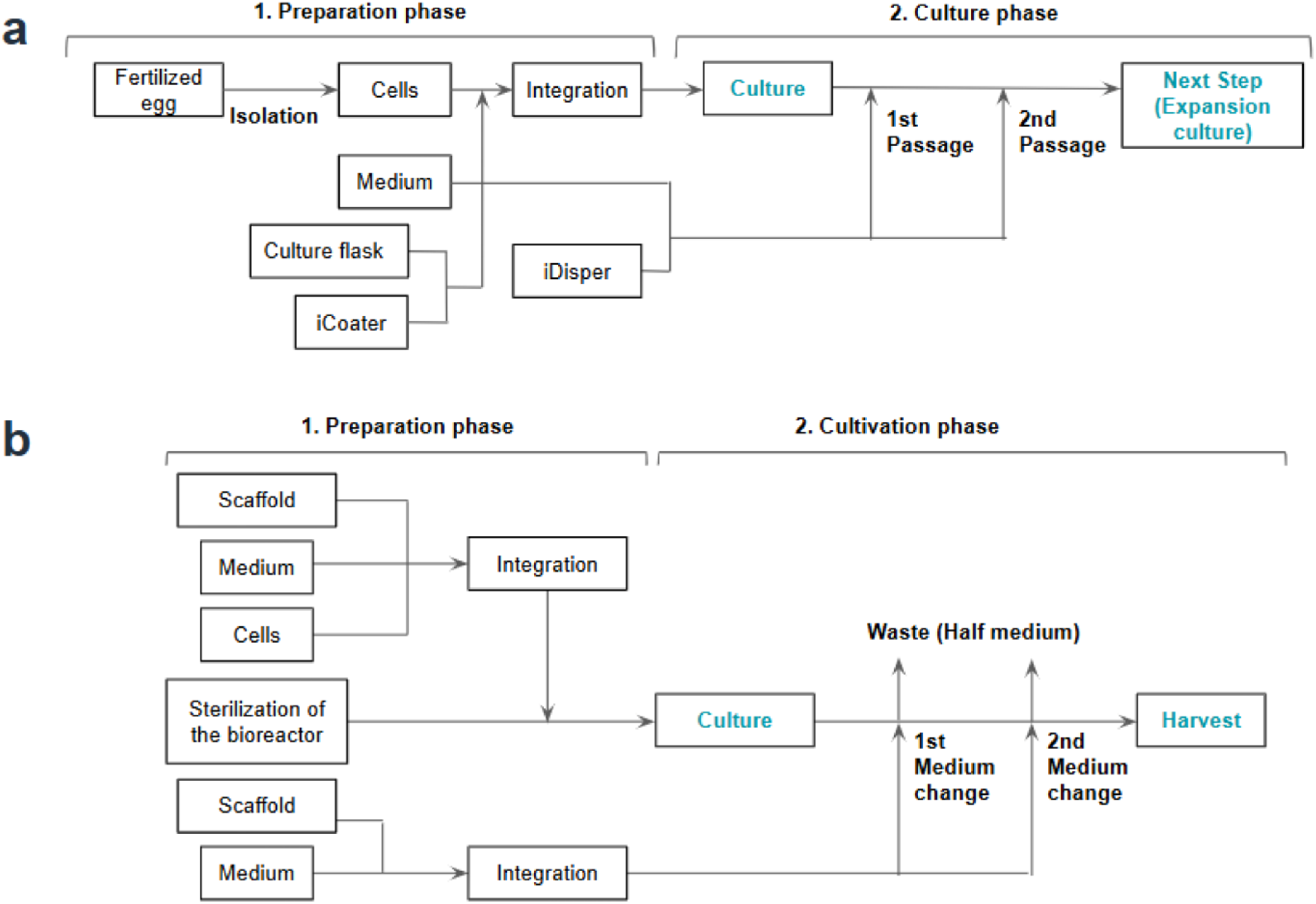
Process flow diagrams of the pre-culture and expansion culture steps. (a) Schematic workflow of the pre-culture step. This process was divided into a preparation phase, involving cell isolation and flask setup, and cultivation phase, where cells underwent two passages to generate a suitable population for expansion. (b) Schematic workflow of the expansion culture step. This process included a preparation phase for bioreactor setup and cell seeding onto scaffolds, followed by a cultivation phase with scheduled medium exchanges, culminating in the final harvest.

**Fig. 3.**
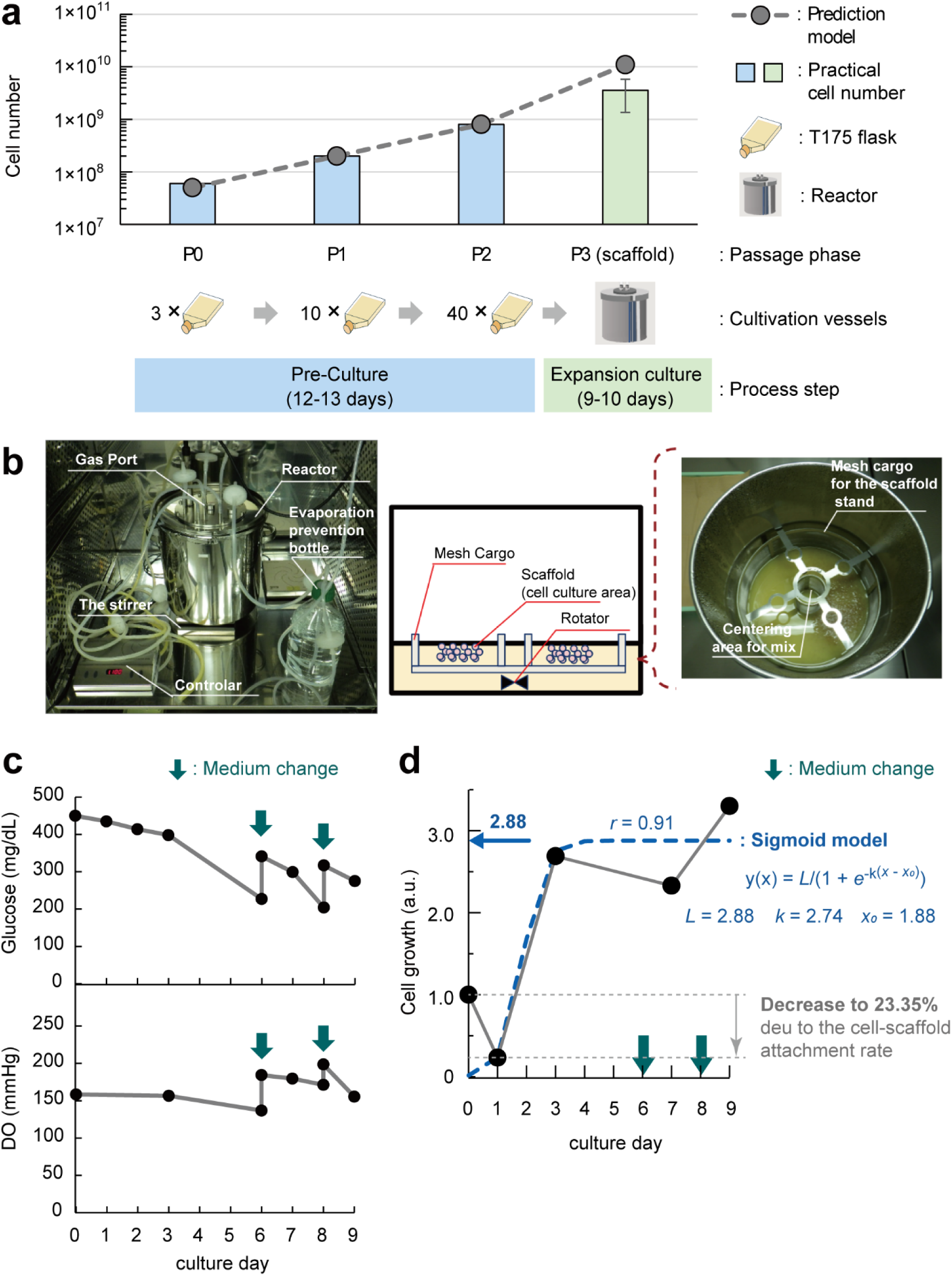
Performance evaluation and scale-up demonstration of the integrated production process. (a) Overall cell number expansion throughout the integrated process. The pre-culture step (12–13 days) involved scaling up cells through passages P0 (3× T175 flasks), P1 (10× T175 flasks), and P2 (40× T175 flasks). The expansion culture step (9–10 days) involved cells at P3 cultured on scaffolds within the bioreactor. The “prediction model” (dashed line) was calculated as described in **Methods S2**. The “practical cell number” (blue bars) represents the average of seven trials (*n* = 7) conducted over a seven-month period in 2025. The variation in process duration (e.g., 12–13 or 9–10 days) was due to operational scheduling, accommodating weekends and holidays. (b) The custom-developed packed-bed bioreactor used for mass production. (c) Time-course analysis of metabolic indicators in the bioreactor. Glucose (mg/dL) and dissolved oxygen (DO; mmHg) levels are shown, with green arrows indicating medium changes. (d) Representative time-course of total viable cell growth (a.u.) during cultivation. The dotted line indicates an initial decrease to 23.35% of the seeded cell number due to the cell–scaffold attachment rate. Green arrows indicate medium changes. The growth followed a “sigmoid model” (*r* = 0.91), shown in blue with fitted parameters (*L* = 2.88; *k* = 2.00; *x_0_* = 2.02), which plateaud at a practical maximum of 2.88-fold.

### 3.3 Performance evaluation: Cell expansion from pre-culture to final harvest

Evaluation of the integrated production process revealed its consistent scalability. Throughout the process, the total cell number increased from the initial cell input of 6 × 10⁷ cells to a final harvest that met the manufacturing acceptance threshold of 3.6 × 10^9^ cells/batch, demonstrating a scale-up from millions to tens of billions of cells (**Fig. 3a**).

This scale-up was founded on the “pre-culture step,” as a stable and scalable production process requires this step to ensure the preparation of a homogeneous cell population before expansion. As shown in **Fig. 3a**, the cell yield in T175 flasks (P0–2) aligned well with the prediction model. As confirmed in our previous report (**Results S1**; Hatano et al., 2025b), this reproducibility was achieved because the initial step selectively amplified a cell population with high proliferative potential, characterized by the increased expression levels of mesenchymal markers compared to those of mature hepatocyte markers.

Such selection occurred because the passaging process, using food-grade reagents, preferentially detached and enriched the cells with high proliferative capacity (mesenchymal or progenitor-like cells), whereas non-proliferative mature cell types were progressively lost. However, **Fig. 3a** also reveals a discrepancy between the theoretical prediction model and practical cell number achieved in the P3 (scaffold) expansion culture, indicating that the full yield potential was not realized in the current 2025 process.

Cell expansion was achieved using the custom-designed packed-bed bioreactor (**Fig. 3b; Fig. S1**). As cells typically consume glucose via aerobic respiration, this was also observed in our system, supported by the steady dissolved oxygen levels, which confirmed sufficient supply in the bioreactor (**Fig. 3c**). Average cell growth exhibited a sigmoid pattern (**Fig. 3d**), which was fitted to a model (*r* = 0.91). This model showed that practical cell growth reached a plateau of approximately 2.88-fold. Quantitative analysis revealed that the practical cell yield (2.88-fold) reached 21.1% of the theoretical maximum (13.65-fold) calculated based on the doubling time (36.8 h) and culture duration (**Methods S2**; Eq. 2).

### 3.4 Process stability and final product safety

The stability of the integrated production process was validated by assessing its reproducibility at critical stages. High concordance was observed in the global gene expression profiles of cells cultured in different manufacturing lots of the food-grade reagent platform (P0: *r* = 0.95; P2: *r* = 0.99). Furthermore, analysis of the same passage-number cells on the scaffolds installed in the bioreactor from different production batches (P_scaffold_ [lot A] vs. P_scaffold_ [lot B]) revealed a high positive correlation between the lots (*r* = 0.98), indicating high (**Fig. 4**). To further assess this quantitatively, we established a 95% prediction interval. The analysis revealed that most genes fell within this narrow interval, providing statistical evidence for the robustness of the manufacturing process. This high degree of consistency is particularly notable because the system used primary cells that are inherently prone to biological variability. The ability to maintain such low lot-to-lot variation underscores the effectiveness of our standardized platform for ensuring product uniformity, a critical attribute for the industrial-scale production of cultivated meat.

**Fig. 4.**
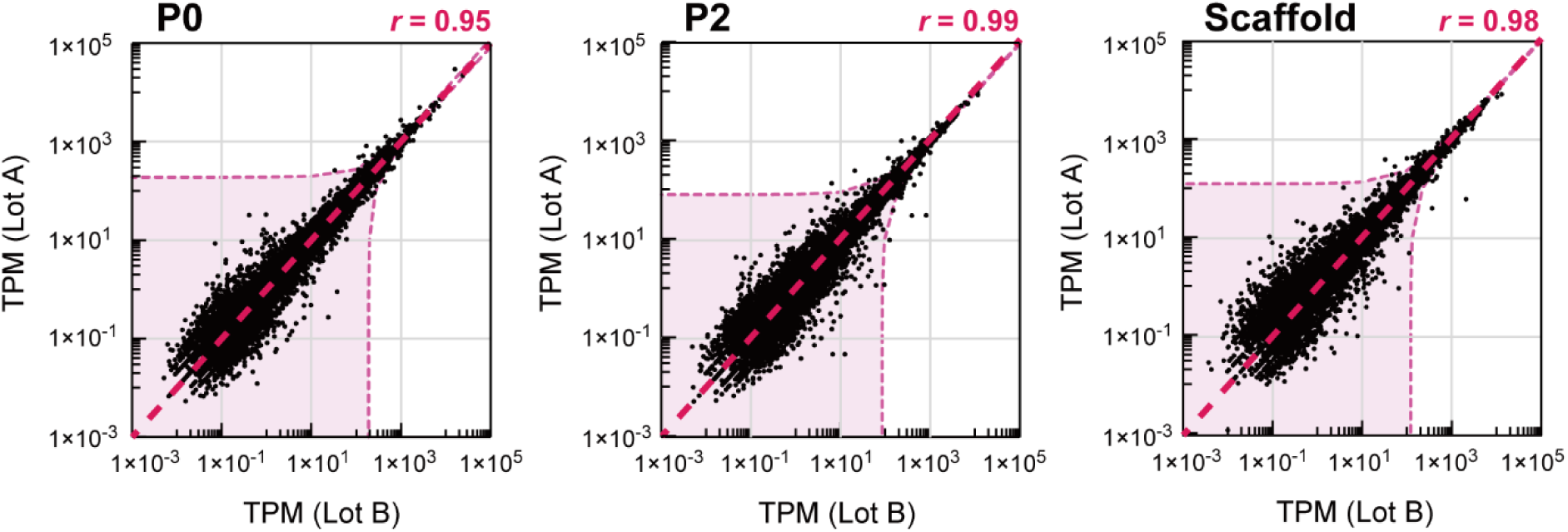
High lot-to-lot reproducibility of gene expression. Assessment of process reproducibility. High concordance in global gene expression profiles was observed between cells cultured in different manufacturing lots (lots A and B) of the food-grade reagent platform (left, P0 and P2) and those cultured on the scaffold (right). The light pink shaded area delineates the 95% prediction interval, indicating that new data points are expected to fall within this narrow range with high probability. This provides strong quantitative evidence of the high batch-to-batch reproducibility of the standardized process.

Following the harvest, the cell paste with scaffolds was subjected to processing and packaging, including vacuum packaging into 50-g pouches, heat pasteurization at 75°C for 10–15 min, rapid cooling in ice water, and inspection for metallic foreign objects. The resulting cultivated meat met general food safety standards and was free of microbial and heavy metal contamination (**Fig. 5**). Specifically, it tested negative for harmful pathogens, such as *Salmonella* and coliforms, with concentrations of toxic heavy metals, such as lead and mercury, being well below regulatory limits.

**Fig. 5.**
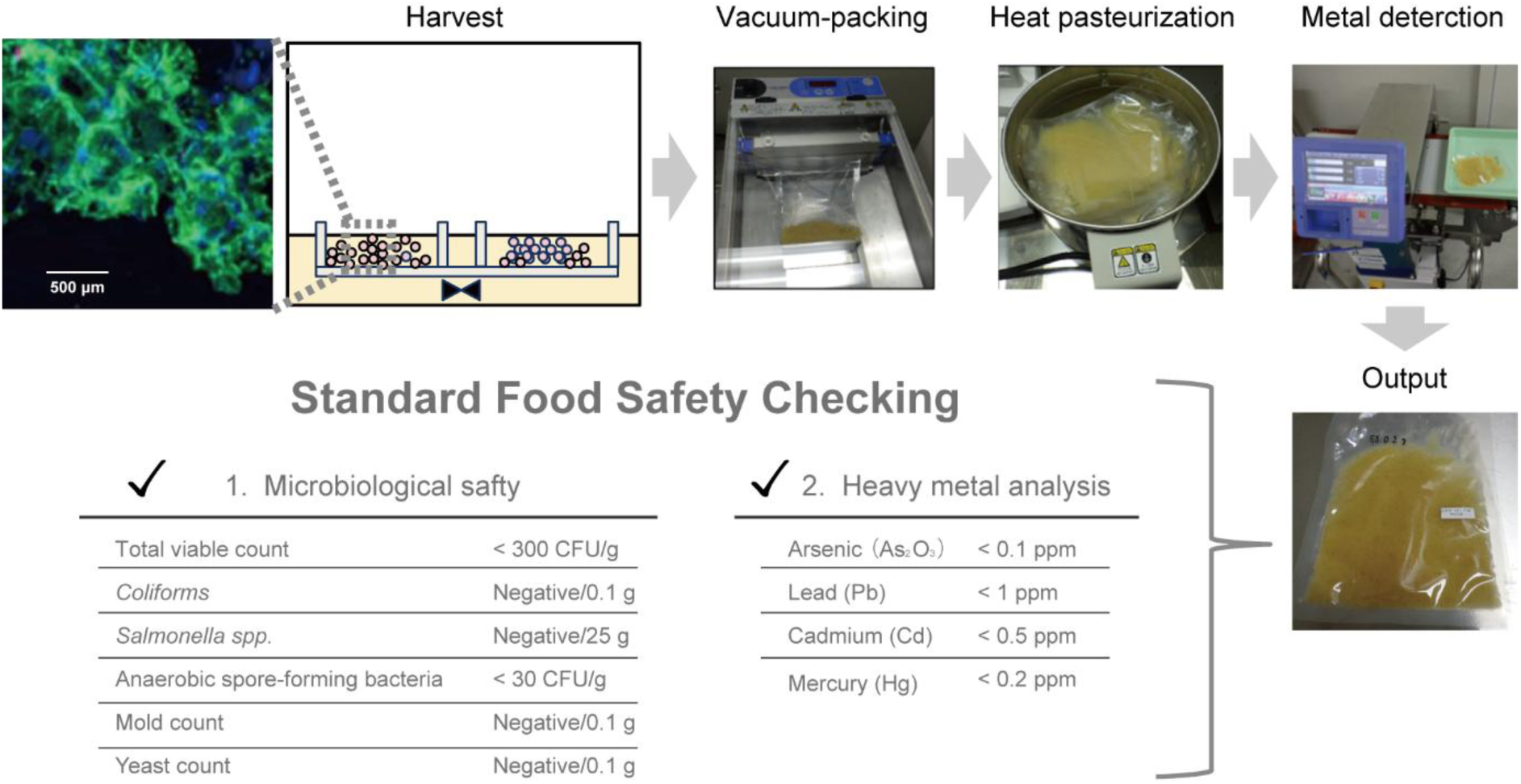
Post-harvest processing and food safety validation of the cell-based food. The process workflow from harvest to the final product packaging, including vacuum packing, heat pasteurization, and metal detection, is depicted. Standard food safety assessments confirmed that the resulting the cell-based food met general safety standards, testing negative for harmful pathogens and showing toxic heavy metal concentrations well below regulatory limits.

Comprehensive nutritional analysis was conducted to characterize the final cell-based product as a food ingredient (**Fig. 6**). Basic analysis confirmed that the product contained significantly fewer calories (energy) than conventional duck liver (**Fig. 6a**). The detailed composition (per 100 g, wet basis) shown in **Fig. 6b** confirmed that the product exhibited high moisture content and low protein and lipid contents on a wet-weight basis. However, on a dry-weight basis, this equated to a high protein content (69.2%). The amino acid composition (**Fig. 6c**) showed a balanced profile, although it exhibited a different pattern compared to that of conventional duck liver, with most amino acids, except for glycine, being present at lower levels. Lipid composition analysis also revealed a distinct low-lipid profile (**Fig. 6d**). Overall, the product ranked significantly lower in all major fatty acid categories, including saturated, monounsaturated, and polyunsaturated fatty acids, compared to the duck liver paste.

**Fig. 6.**
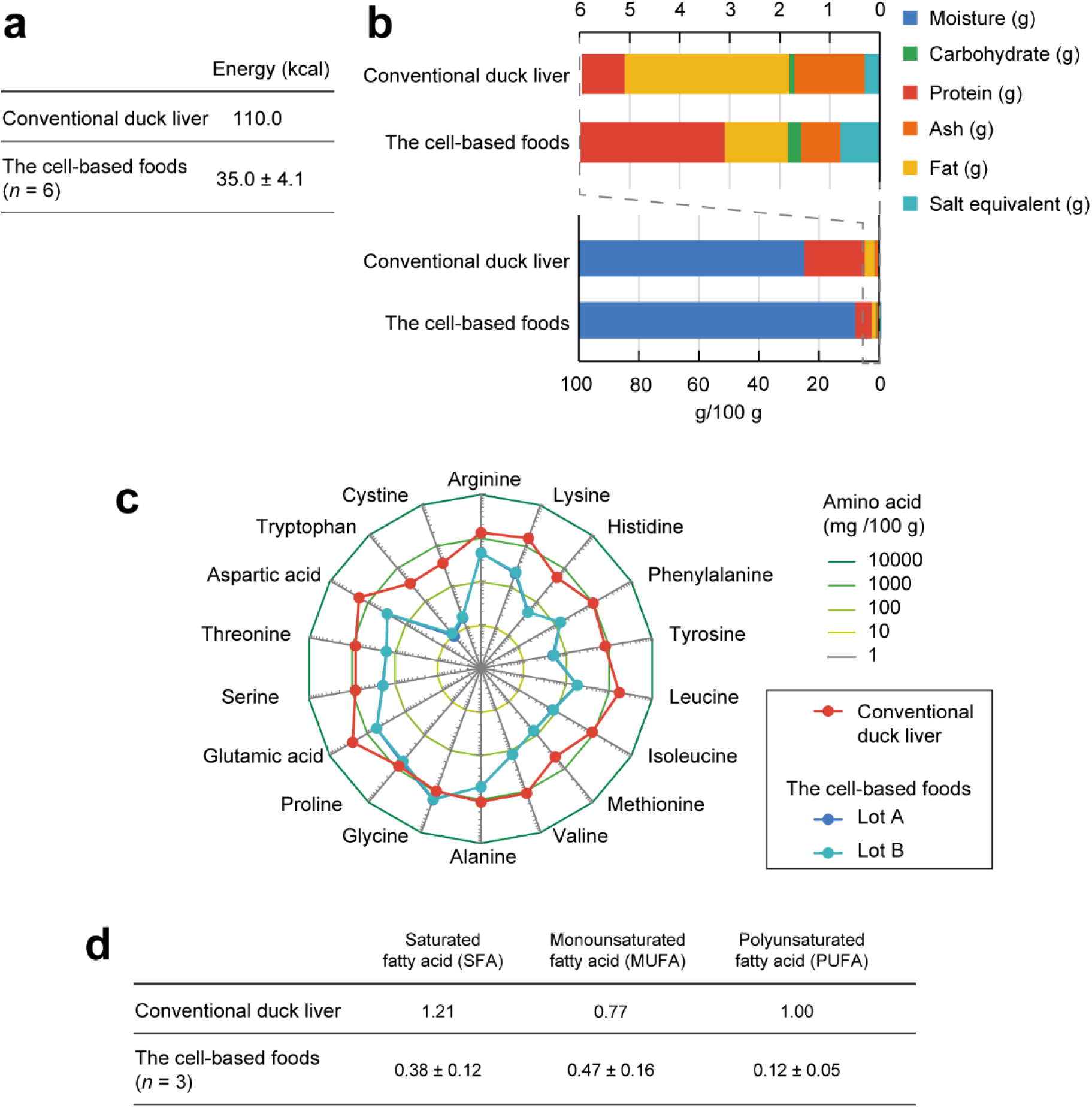
Comprehensive nutritional composition profile of the cell-based food compared to that of conventional duck liver. (a) Energy (kcal) comparison per 100 g (wet basis) with conventional duck liver. (b) Detailed nutritional composition (per 100 g) of the cell-based food, presented as the mean ± standard deviation (SD; *n* = 6 lots): Moisture (92.13 ± 0.62 g), proteins (5.52 ± 0.48 g), lipids (1.3 ± 0.32 g), carbohydrates (0.27 ± 0.18 g), ash (0.78 ± 0.04 g), and salt equivalent (0.78 ± 0.03 g). (c) Amino acid composition radar chart comparing the duck liver paste (red) and cell-based food product (lot A, blue; lot B, teal) in two lots. Notably, the profiles of lots A and B overlapped completely. (d) Fatty acid composition profile (*n* = 3 lots) comparing the cell-based food product with the duck liver paste (g/100 g).

The above-mentioned factors resulted in the distinct “low-lipid, low-calorie” characteristics of the product. To demonstrate its potential, we prepared a novel food prototype incorporating 30% (w/w) of the cultivated meat from this production process (**Fig. S2**). The prototype received high sensory evaluation scores for overall taste and texture (**Table S1**), confirming its culinary viability.

### 3.5 Productivity modeling and future prospects

Finally, to translate our experimental findings into a strategic framework for future scale-up and commercialization, we established a simulation model to estimate the potential monthly production of cell biomass for different animal species. As detailed in **Methods S2**, this model integrated key process parameters, including the intrinsic cell doubling time (*T_d_*), pre-culture expansion folds (*F*_*p*_), and bioreactor culture duration (*D_BR_*). This predictive tool was designed to assess the economic viability of different cell lines and identify bottlenecks in the production process.

Based on the experimentally validated doubling times, comparative analysis was conducted to estimate the monthly production capacity per production unit for each cell line. The calculations revealed that avian-derived cells, particularly duck- and chicken-derived cells, exhibited the highest estimated monthly yields under the established process conditions, whereas mammalian cells showed lower productivity (**Table 1**). These findings suggest that cell type selection significantly affects production efficiency and that avian-derived cells exhibit more cost-effective production profiles than mammalian cells.

**Table 1.**
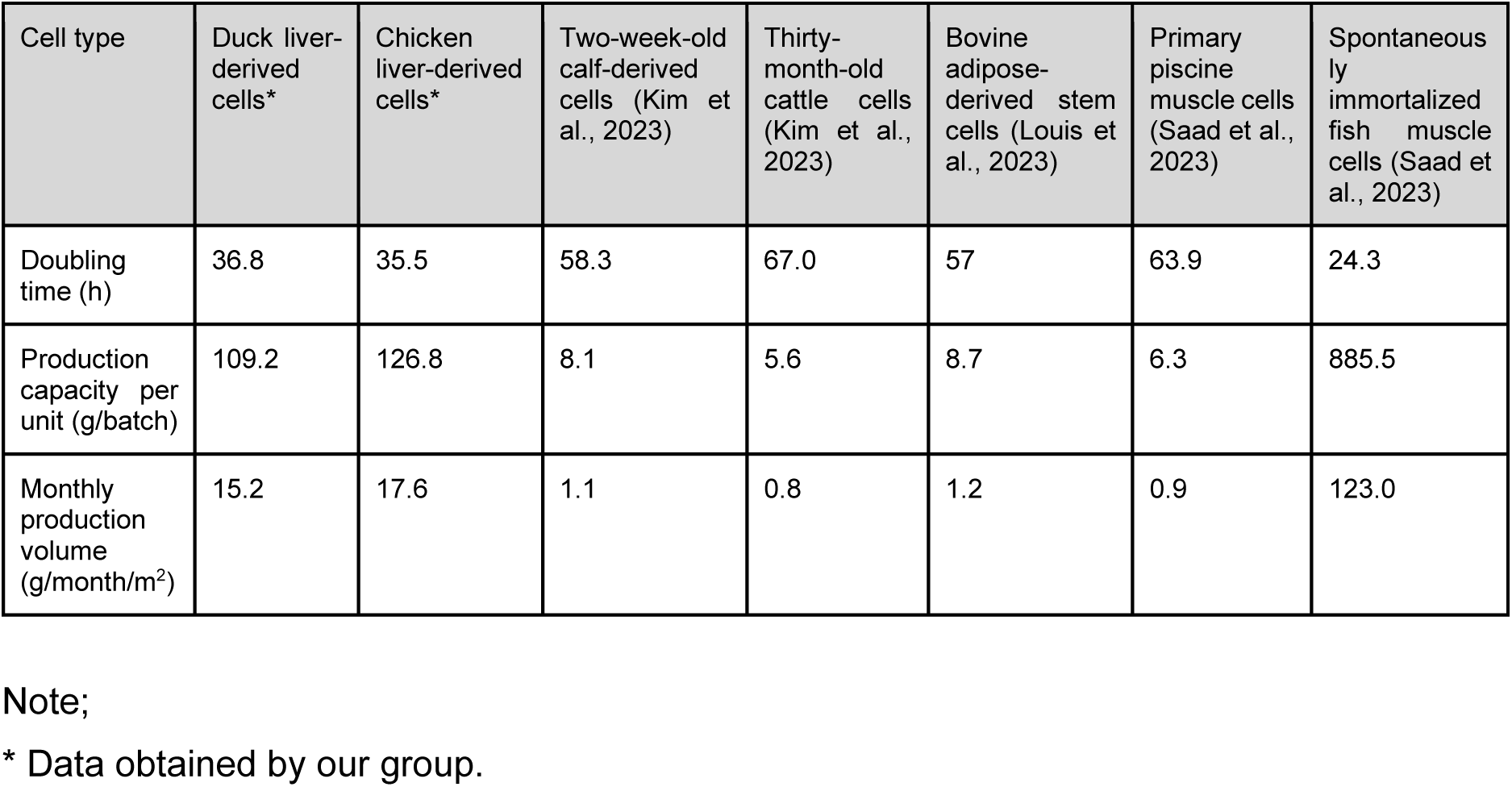
Comparative production performance: Avian cells exhibit high yields in the standardized process.

The above-mentioned estimates provide valuable insights to scale up production and select appropriate cell lines for specific applications. However, these figures are based on a simulation model and can only serve as initial estimates. Further empirical studies are necessary to validate and refine the productivity projections for each cell type.

## 4. Discussion

### 4.1 Addressing the bottlenecks of adherent cell expansion

The primary challenge addressed in this study was the efficient large-scale production of adhesion-dependent cells. Although many current market-approved products (e.g., GOOD Meat and UPSIDE Foods products) use suspension culture processes (Kirsch et al., 2023), adherent cells remain crucial for replicating specific target tissues. Traditional methods, such as agitated cultures with microcarriers, often face limitations in terms of shear stress damage and material costs.

The packed-bed bioreactor system established in this study offers a robust solution to these problems. By providing a three-dimensional scaffold, this system facilitated high-density culture while preserving the phenotype of primary duck liver cells. The success of our approach was evidenced by a 4-fold cell expansion, driven by a stable culture environment that maintained the dissolved oxygen level at approximately 5 ppm to support active respiration.

Importantly, our platform is versatile. Although it was validated using duck liver cells in this study, the scaffold is applicable to various adherent cell types (Hatano et al., 2024), including muscle cells and adipocytes, across different species. Our calculations further suggest that avian-derived cells exhibit higher productivity potential than mammalian cells for this specific process. Future developments should focus on scaling up the working volume, which will require advanced sensing and perfusion systems to maintain environmental consistency at larger scales.

### 4.2 Nutritional profile and market positioning

The cultivated meat produced in this study exhibited a nutritional profile distinct from that of conventional animal products. Notably, the obtained product contained significantly lower levels of saturated fats and total lipids than the conventional duck liver paste, making it a low-calorie food material. These characteristics highlight the potential of the product as a “smart food” for health-conscious consumers or athletes.

The current process results in a product with high moisture content, which lowers the amino acid score on a wet-weight basis. Therefore, future optimizations should focus on media improvement or downstream concentration processes to enhance nutritional density. Moreover, although sensory evaluation confirmed that the texture was favorable, objective instrumental texture analysis in future studies is necessary to correlate sensory scores with rheological properties.

### 4.3 Comprehensive safety assurance

Because cell-based products are classified as “novel foods,” they require rigorous safety evaluations of residual media components and cell by-products (WHO, FAO, 2023). Our process incorporated a heat sterilization step (75°C for 10 min) to reduce these risks by denaturing proteinaceous growth factors and enzymes (Sanchez-Ruiz, 2010; Wolz and Kulozik, 2015).

Heat-stable compounds, such as thyroid hormones and low-molecular-weight peptides, may persist despite sterilization. Therefore, future safety verifications should include:

- **Quantification:** Using high-sensitivity enzyme-linked immunosorbent assay to measure residual growth factor (e.g., insulin-like growth factor-1) and hormone levels.
- **Activity verification:** Testing heat-treated extracts on model cell lines to ensure biological inactivity.
- **Allergen profiling:** Conducting expression profiling for known allergens, such as tropomyosin (WHO et al., 2001).

Finally, we addressed the risk of FBS-associated bovine spongiform encephalopathy (Brown et al., 2001; WHO, 2003, Taylor, 2000). Our FBS-free approach successfully eliminated prion-related risks, stabilized supply, reduced costs, and resolved ethical concerns regarding serum sourcing.

## 5. Conclusion

This study established an integrated cultivated meat production process, including a selective pre-culture step for highly proliferative cells, an expansion culture step to maintain the characteristics of planar culture using a packed-bed bioreactor, and a processing step to obtain the final product, to address the major challenges in cultivated meat production. Through this process, a highly proliferative cell population was stably obtained and cultured. The produced cultivated meat not only met the microbiological and heavy metal safety criteria but also exhibited a unique nutritional profile characterized by low-lipid and low-calorie properties.

Our practical production process also comprehensively verified the safety and quality of the produced meat, overcoming the bottlenecks of mass culturing adherent animal cells. In this process, the packed-bed bioreactor contributed to the efficient expansion and culture of adherent cells. Notably, our production process potentially exhibits high production efficiency, especially for avian-derived cells, and may be applicable to the development of various cell types.

Overall, the production process developed and verified in this study shows potential for producing cultivated meat using adherent cells, thereby providing a novel food material with distinct nutritional characteristics. Through future technological innovation and dialogue with society, safe, delicious, and sustainable cultivated meat will contribute to future food supply and the diversity of food culture.

## Declaration of competing interests

All authors are employees of IntegriCulture Inc. (Kanagawa, Japan). Author Masanobu Kowaka is an employee of both IntegriCulture Inc. and Hamano Products Co., Ltd. (Tokyo, Japan).

IntegriCulture Inc. developed and commercialized the proprietary food-grade reagents (I-MEM 1.0; iDisper; iCoater; IntegriCulture Serum-Replacement) used in this study. Consequently, IntegriCulture Inc. has a direct financial interest in the research findings and their publication and may derive financial benefits from this study. Hamano Products Co., Ltd. designed and supplied the bioreactor (Hamano Reactor v3) used in this study and may have an indirect commercial interest in the reported applications and performance of its equipment.

Beyond the stated affiliations and interests, the authors declare no other known competing financial or personal relationships that could have inappropriately influenced the work reported in this manuscript.

## Declaration of generative AI and AI-assisted technologies in the manuscript preparation process

The authors used Gemini (developed by Google) to improve the readability and language of this article. However, the authors confirm that they have reviewed and edited the content, as required, after using this service and take full responsibility for the content of the published article.

## Data availability statement

The research data supporting the findings of this study are not publicly available because of proprietary and commercial confidentiality. However, the data are available from the corresponding author upon reasonable request.

## Author contributions

**Hiroaki Hatano:** Conceptualization, Methodology, Formal analysis, Writing – Original draft preparation, and Writing – Reviewing and editing. **Ibuki Kokido:** Investigation, Data curation, and Writing – Reviewing and editing. **Keita Tanaka:** Investigation. **Satoshi Inoue:** Investigation, Data curation, and Validation. **Takahiro Sunaga:** Investigation and Validation. **Masanobu Kowaka:** Software and Resources. **Ikko Kawashima:** Supervision, Project administration, and Writing – Reviewing and editing. **Hiroaki Kondo:** Supervision. **Kazuhiro Kunimasa:** Supervision. **Naomi Nakamura:** Resources. **Misaki Sawada:** Resources. All authors have read and approved the final manuscript.

## Funding

This study was supported by the Small Business Innovation Research Program of the Ministry of Agriculture, Forestry and Fisheries of Japan under the “Demonstration of a Production System for Cellular Foods Utilizing CulNet Supernatant” (N020) project. The role of the funding source was limited to financial support.

## Acknowledgements

The authors would like to express their sincere gratitude to all members of IntegriCulture Inc. for their dedicated support and collaboration throughout this study. We are also deeply grateful to Mr. Hideyuki Goto of NPS Inc. (Tokyo, Japan). for his invaluable advice and insightful comments that significantly contributed to this study.

## Glossary

FBS: fetal bovine serum

## Notes

### Competing Interest Statement

The authors have declared no competing interest.

