## Supplementary_information for "An integrated scalable process for adherent cultivated meat production: From proliferative cell selection to safety-verified product development"

**Supplementary Methods**

**S1. Characteristic evaluation of the cell-based food**

To evaluate the characteristics of the cell-based food product, samples of the final processed cell paste (after harvesting, packaging, heating, and freezing, as described in section 2.3 of the main manuscript) were sent for external analysis to the SUNATEC Food Analysis and Development Center General Incorporated Foundation (Mie, Japan). The following analyses were conducted by this institution:

**Microbiological safety profile:** The samples were analyzed for various microbiological indicators to confirm their safety. The tests included total viable counts and detection of coliforms, *Escherichia coli*, *Salmonella* spp., anaerobic spore-forming bacteria, molds, and yeasts, according to the standardized food safety testing protocols. The specific items tested are shown in Fig. 5.

**Heavy metal analysis:** Concentrations of key heavy metals, specifically arsenic (As), lead (Pb), cadmium (Cd), and total mercury (Hg), in the samples were determined using methods such as inductively coupled plasma mass spectrometry. The limits of detection and results are presented in Fig. 5.

**Nutritional composition:** Comprehensive nutritional analysis was conducted. This included the quantification of the moisture, protein, total lipid (fat), carbohydrate, and ash contents. The sodium content was used to calculate the salt equivalent. These analyses provided a proximate composition of the cell-based food product, as shown in Fig. 6.

**S2. Modeling cell production capacity**

A simulation model was established to estimate the potential monthly production of cell biomass per production unit. This model used a common framework based on standard calculations for cell proliferation kinetics and process yields to project the output. It considered key process parameters, including the intrinsic cell doubling time ($T_d$), number of pre-culture passages ($n$), and bioreactor culture conditions. The simulation framework is described below.

**S2.1 Pre-culture yield model**

Passaging was scheduled based on an average interval of 72 h (equivalent to three days). This interval was designed to accommodate operational constraints, such as standard workdays (e.g., passaging on Mondays and Fridays, allowing for 72 ± 8 h between passages). The fold expansion achieved per passage ($F_p$) was dependent on the cell doubling time ($T_d$) and initial seeding density, which was adjusted to ensure that the cells reached near-confluence in the T175 flask:

If $T_d \leq 40$ h: $F_p = 4$ (achieved by seeding at 25% confluence)
If $40 < T_d \leq 50.5$ h: $F_p = 3$ (achieved by seeding at approximately 33% confluence)
If $50.5 < T_d \leq 80$ h: $F_p = 2$ (achieved by seeding at 50% confluence)

Total cell mass after *n* passages ($M_{pre}$) in the preculture step from the initial cell mass $S_0$ was determined using equation (1):

$$M_{pre} = S_0 \times (F_p)^n \qquad (1)$$

#### S2.2 Expansion culture and monthly output model

Following preculture, the cells were transferred to the packed-bed bioreactor for the main expansion phase. The cells were cultured in the bioreactor for a defined period, $D_{BR}$ (typically 9–10 d), and the cell attachment rate of the scaffold ($R_{CA}$) was determined. In this simulation, the cell attachment rate of the scaffold ($R_{CA}$) was predicted to be 23.35% based on our practical data of the current process (2025). As this rate varies with the scaffold material and cell type, empirically measuring this parameter is a key step to refine the predictive accuracy of the model. The fold expansion in the bioreactor ($F_{BR}$) was determined from the cell doubling time ($T_d$) and calculated using equation (2):

$$F_{BR} = 2^{(D_{BR} \times 24)\,/\,T_d} \times R_{CA} \qquad (2)$$

Using the parameters for duck liver cells ($T_d$ = 36.8 h; $D_{BR}$ = 9 d; $R_{CA}$ = 0.2335 [23.35%]), theoretical $F_{BR}$ was calculated as **13.65**. This theoretical value served as the benchmark for the *Prediction model* in Fig. 3a and highlighted the 4.74-fold gap to the *practical* 2.88-fold yield (Fig. 3d) achieved in the current process.

A standard production batch operated $N_{batch}$ bioreactors (e.g., $N_{batch}$ = 4 in our working line in 2025). The total cell mass harvested per batch from one production unit ($M_{unit}$) was calculated using equation (3):

$$M_{unit} = N_{batch} \times M_{pre} \times F_{BR} \qquad (3)$$

The estimated monthly production ($P_M$ in g), based on four harvest cycles per 30-d month in our established process, was determined using equation (4):

$$P_M = M_{unit} \times 4 \qquad (4)$$

### Supplementary Figures

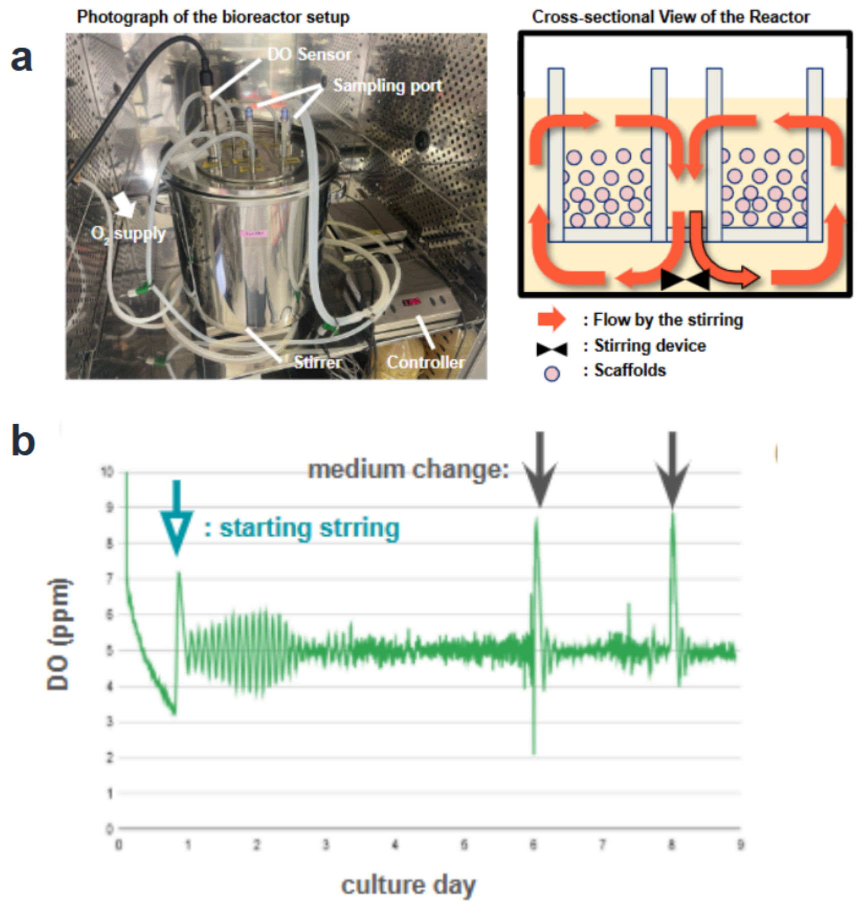


**Fig. S1.** Bioreactor setup and dissolved oxygen (DO) level control. (a) Schematic representation of the 20-L packed-bed bioreactor. The overall experimental setup is shown on the left. The cross-sectional diagram on the right illustrates the internal structure, where the orange arrows indicate the direction of medium flow, ensuring a uniform culture environment. The DO level was maintained at around 5 ppm, the typical level for high-density animal cell culture, via PID-controlled $O_2$ supply, ensuring adequate oxygenation in the bioreactor. (b) DO levels during the 9-d cultivation period. Oxygen was supplied via surface aeration, and DO levels were monitored using the VisiFerm DO ECS 120 H0 sensor (Hamilton) to maintain the target level of 5 ppm. The green arrow indicates the initiation of agitation, and gray arrows indicate the time points of medium change.

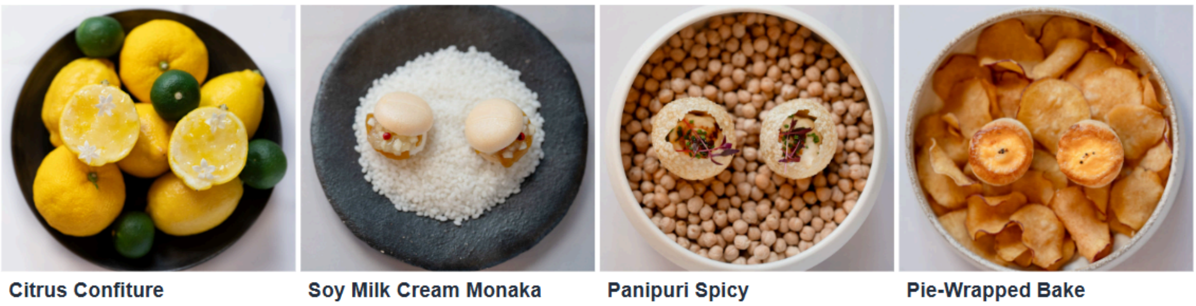

Fig. S2. Prototype "Craft-Essen" menu incorporating approximately 30% the cell-cultivated food obtained via our established production process using duck liver cells. The images were provided by the Craft-Essen Council, an organization in which the authors' institution (IntegriCulture Inc.) is a participating member. The photographs were taken during a sensory evaluation event on February 18, 2025 and are publicly available on the council website (https://craft-essen.com/news-reports/v87d5pj8y3).

**Supplementary Table**

**Table S1.** Summary of the sensory evaluation results for four Craft-Essen menus. Evaluations were conducted by the Craft-Essen Council. The overall taste preference and willingness to try in a restaurant setting were assessed on a 4-point scale and subsequently converted to a 5-point equivalent average score (1 = lowest; 5 = highest; see the calculation method in the following note). Specific sensory attributes (strength and liking of flavor/aroma, liking of texture, strength and liking of aftertaste, and visual appeal) were also rated on a 4-point scale and further converted to a 5-point equivalent average score. The data are presented as average scores (number of respondents, N). Prototypes were developed to explore diverse culinary applications of the cultivated food material.

| Menu name (Prototype) | Brief description | Overall taste | Willingness to try in a restaurant | Flavor strength | Flavor liking | Texture liking | Aftertaste strength |
|---|---|---|---|---|---|---|---|
| Citrus confiture | Sweet and sour dish of cell-based food and soy milk cream stuffed into citrus fruits, accented with rock salt. | 4.7 (N = 29) | 4.1 (N = 30) | 4.3 (N = 31) | 4.0 (N = 31) | 4.4 (N = 31) | 4.0 (N = 31) |
| Soy milk cream Monaka | A dish of spore cell-based food, soy milk cream, pear, jam, pink pepper, and Nara pickles stuffed in the middle. | 4.5 (N = 31) | 4.1 (N = 31) | 3.8 (N = 31) | 4.1 (N = 31) | 4.4 (N = 31) | 3.8 (N = 31) |
| Pani puri spicy | Indian snack “Pani puri” served in a bowl with cell-based food, along with spices and flowers. | 4.6 (N = 31) | 4.2 (N = 31) | 4.3 (N = 31) | 4.3 (N = 31) | 4.2 (N = 31) | 4.0 (N = 31) |
| Pie-wrapped bake | Cell-based food and sweet potato wrapped in pie crust and baked. | 4.2 (N = 30) | 4.0 (N = 30) | 3.8 (N = 31) | 4.0 (N = 31) | 4.1 (N = 31) | 3.8 (N = 31) |

Note:

Average scores were calculated based on a 4-point rating scale (e.g., overall taste: 1 = bad, 2 = not very good, 3 = okay, and 4 = delicious) and subsequently converted to a 5-point equivalent scale using the following formula: (average score on the 4-point scale/4) × 5.
For example, 3.72 points/4 × 5 = 4.65 ≈ 4.7 points.

The number of respondents (N) varied slightly for some attributes due to occasional missing responses, as noted in the source document. “Key positive comments” and “Key areas of Improvement” are summarized from the qualitative feedback provided by the respondents in the source document.
